# Non-invasive and unbiased assessment of thermogenesis in mice through thermal gradient ring

**DOI:** 10.1101/2025.02.17.638667

**Authors:** Paulo De Melo, Nayara Pereira, Rafaela Braun Araujo, William T. Festuccia, Thiago Mattar Cunha, Luiz Osório Leiria

**Author notes:** **Corresponding Author:** Luiz Osório Silveira Leiria, PhD Department of Pharmacology Department of Cell Biology Ribeirão Preto Medical School University of São Paulo Ribeirão Preto, SP - 14049-900.

## Abstract

Accurately assessing whole-body heat production requires reliable thermometry methods. In mice, common approaches include rectal temperature (RT) measurement, infrared (IR) thermography, and implanted probes. However, factors such as stress, handling, surgery, and variability limit their applicability for evaluating thermogenesis. The Thermal Gradient Ring (TGR), widely used in neuropathic pain and ion channel studies, consists of a circular structure with twelve temperature zones and an integrated IR camera for real-time behavior monitoring. This system allows precise analysis of preferred temperature (PT), heat tolerance, locomotion, and zone occupancy over time, thereby offering a behavioral perspective beyond traditional thermometric methods, which provides only temperature data. In this study, we evaluated TGR as a non-invasive tool for detecting thermogenic changes. Since mice with higher thermogenesis prefer cooler zones, while those with reduced thermogenesis seek warmth, TGR provides a sensitive readout of metabolic behavior. Using models with both enhanced and impaired thermogenesis, we demonstrated TGR’s ability to detect thermogenic status under different conditions. These findings suggest that TGR is a valuable tool for metabolic research, offering a reliable alternative for assessing thermogenesis in mice.

## INTRODUCTION

Thermoregulation is a fundamental physiological process in mammals, maintaining stable internal body temperature despite environmental fluctuations (Cannon & Nedergaard, 2011; Glass et al., 2021; Ivanov, 2006). It is essential for cellular function, enzymatic activity, and overall metabolic homeostasis. Mammals regulate body temperature through both behavioral and physiological mechanisms, including seeking warmer or cooler environments, adjusting blood flow, and modulating heat production via shivering and non-shivering thermogenesis (NST) (Ivanov, 2006; Cannon & Nedergaard, 2011; Hymczak et al., 2021). Brown adipose tissue (BAT) plays a central role in NST, generating heat through uncoupling protein 1 (UCP-1) activation and futile cycles. Sympathetic β-adrenergic stimulation drives NST, with BAT-derived heat rapidly increasing core temperature (Cannon & Nedergaard, 2004; Cypess et al., 2009). Beyond cold adaptation, thermoregulation is closely linked to energy metabolism, influencing food intake, body composition, and metabolic rate. Dysregulated thermogenesis is associated with metabolic disorders such as obesity and cachexia, underscoring its significance in overall health (Cypess et al., 2009; Ono-Moore et al., 2020; Petruzzelli et al., 2014; Stanford et al., 2013). Investigating thermoregulation provides critical insights into energy balance and metabolic adaptations, making it a key research focus in physiology and medicine.

In addition to calorimetric essays, accurate thermometry techniques are essential for phenotyping thermogenesis in mice. Common methods include RT measurement, implanted probes, and IR thermography, each with advantages and limitations. Rectal thermometry is simple, affordable, and widely used but is invasive and prone to variability due to probe insertion depth and handling stress (Antonacci et al., 2019; Zethof et al., 1994). Implanted probes enable wireless temperature monitoring in freely moving mice and are valuable for longitudinal studies, yet they require surgery, anesthesia, and may trigger immune responses that affect thermoregulation (Mei et al., 2018). IR thermography provides a non-invasive assessment of surface temperature but is highly variable due to factors such as posture and fur coverage. While shaving can improve accuracy, it may also irritate the skin and alter thermoregulatory responses (Meyer et al., 2017; Ujisawa et al., 2024). Given these limitations, more precise and integrative thermometry solutions are needed in metabolic research.

TGR system offers a novel approach to thermotaxis analysis in rodents. Featuring a circular running track segmented into 12 distinct temperature zones (ranging from warm to cool), the TGR system allows free movement while an IR camera continuously tracks and records the animal’s behavior (Fig. 1). This design eliminates spatial cues and border bias inherent to linear systems, enhances accuracy through duplicate measurements, removes edge artifacts, and automates data acquisition, reducing observer bias and subjective interpretation (Touska et al., 2016). The TGR has been applied in studies of neuropathic pain and thermal sensory mechanisms in peripheral neurons (Touska et al., 2016; Ujisawa et al., 2024).

**Figure 1:**
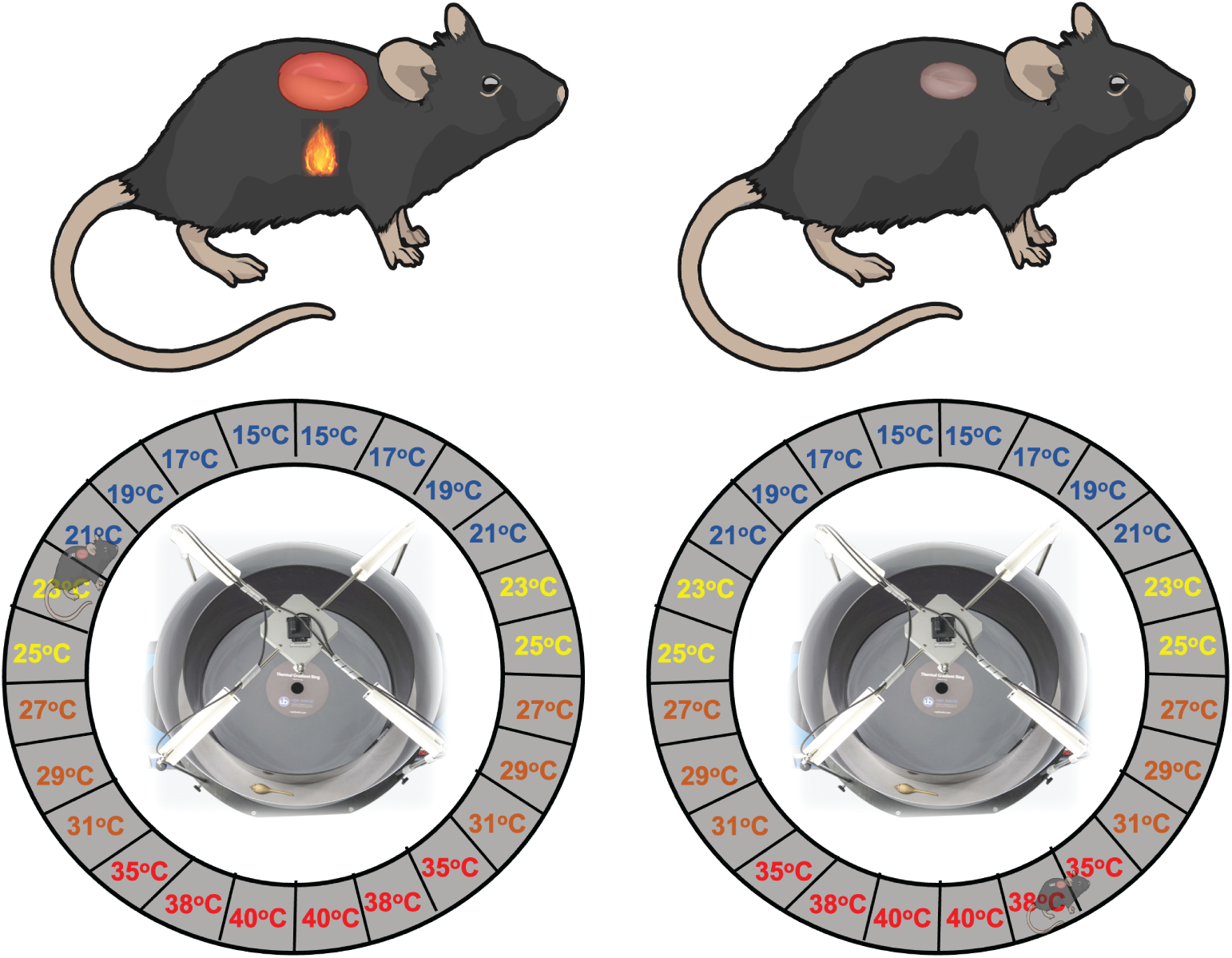
Working hypothesis for thermogenesis modulation using the TGR system. The TGR apparatus comprises a circular platform equipped with twelve-temperature-zone gradient system, enabling real-time behavioral monitoring along a circular track via an IR camera. This system is specifically designed to evaluate temperature-dependent behaviors, including PT, thermal tolerance, locomotor activity, and spatial occupancy over time. By modulating thermogenesis through gain- and loss-of-function interventions in BAT, we hypothesized that mice with enhanced BAT-mediated thermogenesis would exhibit a preference for cooler zones, whereas counterparts with impaired thermogenesis would display a greater propensity for warmer regions.

This study explores the potential of the TGR system for assessing thermogenesis in mice, offering a behavioral perspective beyond traditional thermometric methods, which solely provide temperature data (Lei et al., 2023; Sasajima et al., 2022). The applicability of this method is based on the principle that mice with higher thermogenic capacity tend to seek cooler temperatures, whereas those with lower thermogenesis prefer warmer environments. Using models of BAT gain and loss of function—including β-adrenergic activation, high-fat diet feeding, BAT lipectomy, UCP-1 deficiency, and cold challenge—we demonstrate that TGR is a highly sensitive, non-invasive tool for quantifying thermogenic status. Notably, TGR-detected changes in PT were more sensitive than temperature data obtained via IR thermography or rectal measurements. These findings suggest that TGR can provide high-resolution, reproducible data, integrating both metabolic and behavioral components of thermoregulation in mice.

## METHODS

### Animals

Mice experimental procedures were approved by the Animal Welfare Committee of the Ribeirão Preto Medical School, University of São Paulo (CEUA protocol number: 1196/23). All mice used were males of a C57BL/6J background from Jackson Laboratories (Bar Harbor, Maine, USA). UCP-1 knockout (KO) mice and littermate wild-type (WT) mice were generated by crossing heterozygous UCP-1 KO mice (B6.129-Ucp1tm1Kz/J, Jackson Laboratories, Bar Harbor, Maine, USA). UCP-1 KO mice were kindly bred, genotyped, and provided by Dr. Willian T. Festuccia laboratory. All mice were maintained under specific pathogen-free conditions at the Animal Facility of the Ribeirão Preto Medical School, University of São Paulo. (USP). All experiments were conducted with 7-9-week-old male mice according to the guidelines of the Animal Welfare Committee. Mice were kept at 23 ± 1°C on a 12/12 h light-dark cycle (7 AM/7 PM) and fed a chow diet (70% carbohydrate, 20% protein, 10% fat, in % kcal) or a high-fat diet (20% carbohydrate, 20% protein, 60% fat, in % kcal, with lard as the fat source, HFD).

### Thermal gradient ring (TGR) apparatus and software

The TGR apparatus (Ugo Basile) and software were detailed in Touska et al. (2016). All mice were acclimated for 30 minutes one day before the first run on a device without a thermal gradient at an average ambient temperature of 23°C. The experimental runs were conducted between 08:00 and 17:00 h, maintaining the same schedule for each mouse in both trials. The apparatus used was a small 40 cm diameter aluminum ring-shaped disk, 1.5 cm thick, which allows the mouse to move freely. The inner walls, made of Plexiglas, measure 88 cm, while the outer wall, made of aluminum, measures 141 cm, both with a height of 12 cm. The circular surface area of 640 cm² has a width of 6 cm, where a temperature gradient is equilibrated. This surface provides enough contrast to detect the mouse and allows for surface temperature control using an IR camera (C920, Logitech) located on top of the apparatus. We conducted a 60-minute run, during which the running track and mouse behavior were recorded with a regular CCD camera. The surface temperature was monitored by the IR camera, and all minor deviations (less than 1°C) were readjusted. There are 12 zones of different temperatures (15-40°C) within an interval of 2-3°C from each zone, displayed symmetrically to yield two semi-circles of even temperature on opposite sides. For each temperature zone, the recorded values are provided in duplicate, referring to the opposite side equal temperature zone. The software detection algorithm responsible for automatic data analysis was also previously described(Lei *et al*, 2023; Sasajima *et al*, 2022). In summary, the algorithm detects each mouse position in a circular device in real-time (frame-by-frame) using a computer, so the mouse is assigned to a zone in each frame. This tracking system recorded the number of times the mouse entered each zone, the duration spent in each region, and the overall movement pattern throughout the trial. The data were processed using ANY-Maze software (Stoelting), which facilitated the analysis of various behavioral metrics related to the animal’s movement within specific zones. These metrics included “Time in a Zone,” which refers to the duration an animal spends within a zone, measured from entry to exit in seconds. “Zone Occupancy” was calculated as the percentage of time spent in each zone over the course of the experiment. The “Cumulative Distance from the Zone” was defined as the product of the distance from the zone and the time the animal spent at that distance, measured in meters multiplied by seconds. “Preferred Temperature (PT)” was determined as the average temperature of the zone the animal visited over time. The “Distance Covered” metric represents the total distance traveled across all sequences that ended within the specified time period, measured in meters. Finally, the “Average Time Spent in the Zone” was calculated as the mean duration the animal spent in the chosen zone, with a focus on zones above 35°C. To further illustrate the animal’s behavior, a heat map was generated, providing a visual representation of the mouse’s zone preferences and locomotion patterns within the TGR system.

### iBAT lipectomy (iBATx) surgery

As previously described (Morriseyt & Carnie, 1982) C57BL/6J young male mice (8 weeks old) underwent bilateral iBATx. Briefly, the mice were anesthetized with inhalation administration of isoflurane, which was maintained throughout the surgery. The procedure was conducted in an aseptic manner. Before the incision, the hair around the interscapular area below the neck was shaved, and the mice were placed in ventral recumbency. Following preparation, a 2-cm incision was made along the dorsal midline, and bilateral pads of BAT were removed along with the surrounding white adipose tissue. The area was cauterized, and a midline suture was used to close the incision. Dipironne at a dosage of 80 mg/kg was administered via intraperitoneal injection immediately after surgery. Daily supervision was conducted until day 4, when all mice showed complete recovery of body weight. The TGR analysis was performed on day 4 post-lipectomy.

### CL316,243 injection

Immediately after a first run, each mouse had their thermographic picture taken, the weight and RT measured and received a subcutaneous injection of CL316,243 (1 mg/ kg, ID: 24278325 Sigma-Aldrich, C5976) near the interscapular brown adipose tissue (IntraBAT). Body temperature was monitored until an elevation of at least 0.5°C was observed before initiating the second run of TGR.

### Cold exposure

After the first run in TGR, 8-week-old male mice were placed in individual cages at cold temperatures (4-8°C) for 4 days. The cold chamber was ventilated with a 12-hour light/dark cycle (lights on from 07:00 to 19:00 h). Food, water, and cages were changed every 2 days, and the mice’s health was checked daily. On day 4, at the same time as in the first run, mice were individually removed from the cold chamber and placed at 22-24°C for one hour before the second TGR test run. During this time, the thermographic image, weight, and RT were measured.

### High-fat diet feeding

Mice were maintained on a high-fat diet (HFD) for 7 days. On day 0, before initiating the HFD, mice underwent thermographic imaging, were weighed, and had their RT measured. Subsequently, they were introduced into the TGR for analysis. On days 3 and 7, mice underwent thermographic imaging, were weighed, and had their RT measured. Subsequently, they were introduced into the TGR for analysis. HFD composition: Casein, dextrinized starch, lard, L-cystine, choline bitartrate, soybean oil, biphasic calcium phosphate, cellulose, AIN-93M mineral mix, AIN-93 vitamin mix, sucrose, TBHQ (tertiary butylhydroquinone).

### Thermographic images

Mice were placed in a specialized plastic apparatus designed for individual animal thermographic photography. A thermal camera (FLIR E6xt) captured multiple images to accurately assess the animal’s temperatures. These thermographic images were then analyzed using the FLIR Tools application (Franco *et al*, 2019; Vogel *et al*, 2016).

### Rectal temperature

Mice were immobilized by the operator hand and a thermometer (BAT 10 - physitemp instruments LLC) coated with vaselin was gently inserted into the rectum to minimize discomfort. The temperature was allowed to stabilize to ensure the recorded data was as accurate as possible.

### Statistical analysis

Statistical analyses were performed considering a significance level of 0.05 (α = 5%). All analyses were conducted using Spyder (Python 3.12). Statistical tests were performed with the pingouin package (version 0.5.5), while scipy (version 1.13.1) and statsmodels (version 0.14.2) were used for additional statistical analyses, including correlation tests, confidence intervals, and regression modeling. The scikit-learn package (version 1.5.1) was employed for data preprocessing and specific machine learning analyses. Additionally, pandas (version 2.0.3) and numpy (version 1.25.0) were used for data organization and numerical computations. To ensure statistical robustness, data that did not adhere to normality were transformed using the Box-Cox method. Outlier exclusion was strictly limited to measurement or technical errors. Assumptions of normality, linearity, homoscedasticity, and multicollinearity were assessed when required for the analyses. All graphs and plots were generated using the matplotlib (version 3.9.2) and seaborn (version 0.13.2) packages. All scripts and results were documented to ensure reproducibility and traceability.

## RESULTS

### β3 adrenergic receptor (β3-AR) activation leads to a decrease in preferred temperature

Activation of β3-AR promotes lipolysis and UCP-1-dependent and independent thermogenesis in white and brown fat (Kim *et al*, 2022; Feldmann *et al*, 2009; Okamatsu-Ogura *et al*, 2007). Increases in core body temperature (CBT) can be detected around 10 minutes after β3-AR agonist (CL316,243) injection (Kim *et al*, 2022). Here, we investigated whether the increase in CBT (metabolic thermogenesis) induced by CL316,243 results in changes in mice comfortable temperature and whether the TGR is an accurate method to measure this phenomenon. C57BL/6J young adult male mice were individually placed in the TGR ring floor for 60 minutes before CL316,243 injection (Figure 2A). The absolute time spent in the 12 temperature zones were measured through TGR analysis, which revealed that CL316,243 treatment promoted a shift towards the colder zones (Figures 2B). TGR can also provide comprehensive and representative images of mice running through the temperature zones. The heat map shows the preferred temperature zones over 60 minutes of testing; before CL316,243 injection, mice typically stopped in zones around 29°C-35°C, while after CL316,243 injection, the stopping zones ranged between 25°C-29°C (Figure 2C). Also, the cumulative distance from the zone plot indicates that CL316,243-treated mice prefer cooler than warmer zones (Figure 2D). CL316,243 injected mice exhibited lower heat tolerance, as evidenced by their preference for temperatures below 35°C (Figure 2E), and displayed lower mobility (Figure 2F). Moreover, CL316,243 injection raises the rectal temperature by approximately 1°C, which was in contrast with a reduction of about 2.5°C in PT (Figure 2G-I), suggesting that TGR is a sensitive method for evaluating preferred temperature in the context of metabolic thermogenesis. Moreover, we also employed IR thermography analysis to confirm the changes in mice temperature, showing a similar pattern to rectal temperature (Figure 2J and L). Together, these data demonstrate that the TGR is a sensitive tool for evaluating temperature-associated behaviors in a sympathetic activation model.

**Figure 2:**
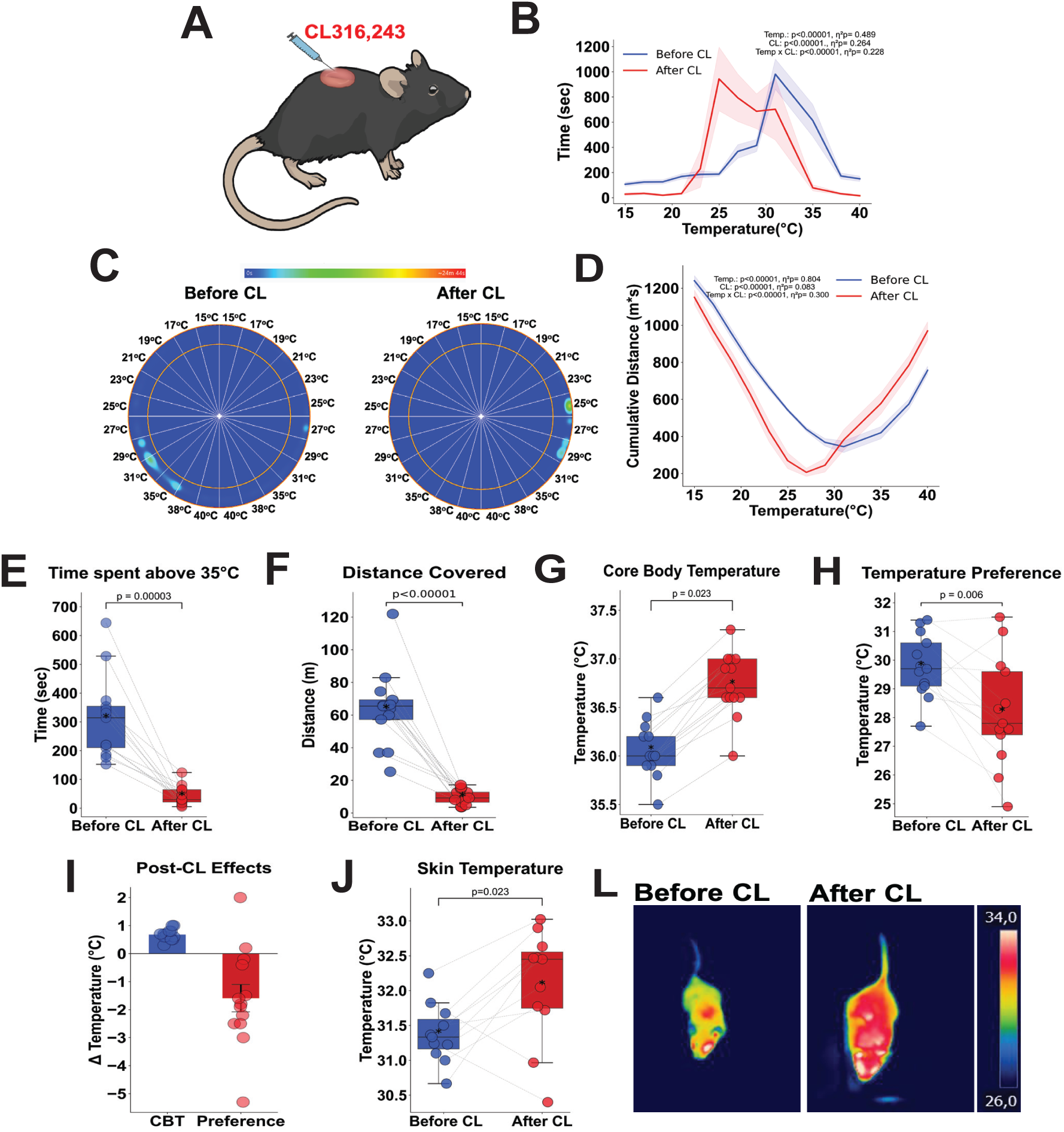
CL316,243 administration increases core body temperature and modulates cold zone preference in mice. Eight-week-old male C57BL/6 mice (n = 10) were evaluated in TGR system before and 20 minutes after intra-BAT injection of CL316,243 (1 mg/kg body weight). (A) Schematic of CL316,243 administration protocol. (B)Time spent per temperature zone within the TGR system during a 60-minute observation period. (C) Representative thermal occupancy heatmap depicting murine locomotor patterns in the TGR. (D) Cumulative distance from the zone across experimental groups. (E) Time spent in temperature zones over 35°C in a 60-minute interval. (F) Locomotor activity during TGR assessment. (G) Rectal temperature measurements before- and after intervention. (H) Mean PT within the TGR system. (I) Comparative analysis of post-intervention Δ changes in CBT and PT. (J) Skin temperature proximal to BAT depots. (L) Representative infrared thermograms of BAT-associated area. Data are presented as mean ± SEM; p < 0.05 (α = 5%). Statistical analysis: Two-way repeated-measures ANOVA (panels B-D); paired t-test (panels F–L).

### Assessment of diet induced thermogenesis by TGR

Dietary fat increases the metabolic rate and triggers thermogenesis (Kinabo & Durnin, 1990; Ono-Moore *et al*, 2020; Nedergaard & Cannon, 2022). A high lipid content in the diet is associated with an increase in UCP1-dependent thermogenesis in (Aquila *et al*, 1985; Fromme & Klingenspor, 2011). Next, we tested whether TGR can be an appropriate method to assess diet induced thermogenesis in mice fed with a HFD for 1 week. C57BL/6J young adult male mice were individually tested in the TGR system before and after one week exposure to HFD (Figure 3A). Notably, mice under a HFD spent longer time at cooler zones (Figures 3B). The heat map illustrates the preferred temperature zones over 60 minutes of testing; before HFD, the mice typically stopped in zones around 28°C-35°C, while after one week of HFD, the stopping zones were between 19°C- 25°C (Figure 3C). Additionally, the cumulative distance from the zone plot indicates a shift from warmer to cooler zones over short term HFD feeding (Figure 3D). Indeed, the time spent above 35°C was shorter after HFD intake (Figure 3E), suggesting a warmth avoidance behavior. HFD intake did not modify mice mobility (Figure 3F), while promoting an increase in CBT (Figure 3G) and in the preference for lower temperatures in TGR (Figure 3H). We found a reduction of 3.5°C in PT, whereas the average increase in CBT was about 1.5°C (Figure 3I). Thermographic images also show an increase in superficial temperature after short-term HFD (Figure 3J and L), which also validates the TGR data. Those data show that indeed TGR is adequate for assessing thermogenesis in thermogenic phenotypes such as induced by sympathomimetic or dietary stimuli.

**Figure 3:**
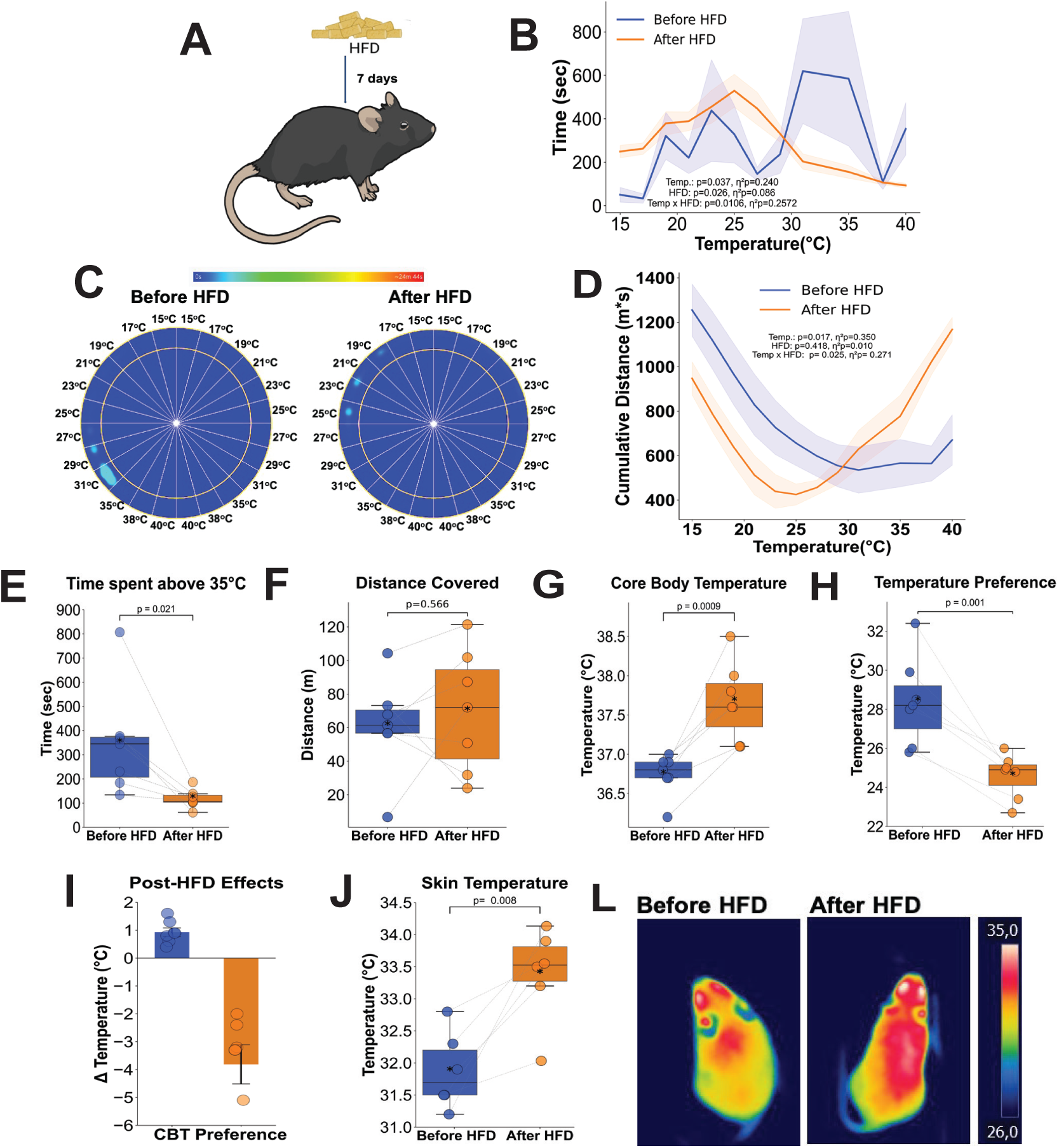
HFD increases core body temperature and shifts thermal preference toward colder zones in mice. Six-week- old male C57BL/6 mice (n = 7) were evaluated in a TGR system before and after 7 days of HFD intervention. (A) Schematic of experimental HFD administration protocol. (B) Time spent per temperature zone within the TGR system during a 60- minute observation period. (C) Representative thermal occupancy heatmap depicting murine locomotor patterns in the TGR. (D) Cumulative distance from the zones across experimental groups. (E) Time spent in temperature zones above 35°C in 60- minute interval. (F) Locomotor activity profiles during TGR assessment. (G) Rectal temperature measurements before and after HFD. (H) Mean of PT within the TGR system. (I) Comparative analysis of after HFD Δ changes in CBT and PT. (J) Skin temperature proximal to BAT depots. (L) Representative infrared thermograms of BAT-associated areas. Data are presented as mean ± SEM; p < 0.05 (α = 5%). Statistical analysis: Two-way repeated-measures ANOVA (panels B-D); paired t-test (panels E-J).

### Interscapular BAT removal alters mice preference for warmer zones

Interscapular BAT surgery removal (lipectomy, iBATx) has been employed as a tool to assess the relevance of BAT in various aspects of metabolism and thermogenesis (Grunewald *et al*, 2018; Jia *et al*, 2021; Morriseyt & Carnie, 1982). Lipectomy of BAT reduces body temperature a few days after surgery, while it is restored to normal levels three weeks post-lipectomy, which may be due to compensatory mechanisms such as the browning of white adipose tissue (Grunewald *et al*, 2018; Jia *et al*, 2021). We sought to investigate whether the low thermogenic capacity typical of the initial days post lipectomy could be sensibly detected through TGR. We performed the TGR test before and after the lipectomy in young adult C57BL/6 male mice (Figure 4A). Seven days after lipectomy, iBATx mice showed a shift in the preference towards the warmer zones (Figures 4B), which can also be observed in the heat map and through the cumulative distance from the zone plot (Figures 4C and D). Additionally, iBATx mice spent more time above 35°C (Figure 4E) and (Figure 4G), in contrast to increased PT (Figure 4H) after surgery. BAT lipectomy promoted an average increase in PT of 2.5°C, while the average reduction in CBT was around 1°C (Figure 4I), once more highlighting the higher sensitivity of this method. Thermographic images confirmed the reduction of superficial temperature in the interscapular BAT area after first weeks of lipectomy (Figure 4J and L). These data indicate that the lower thermogenic capacity resultant from iBAT removal is sensibly detected by TGR.

**Figure 4:**
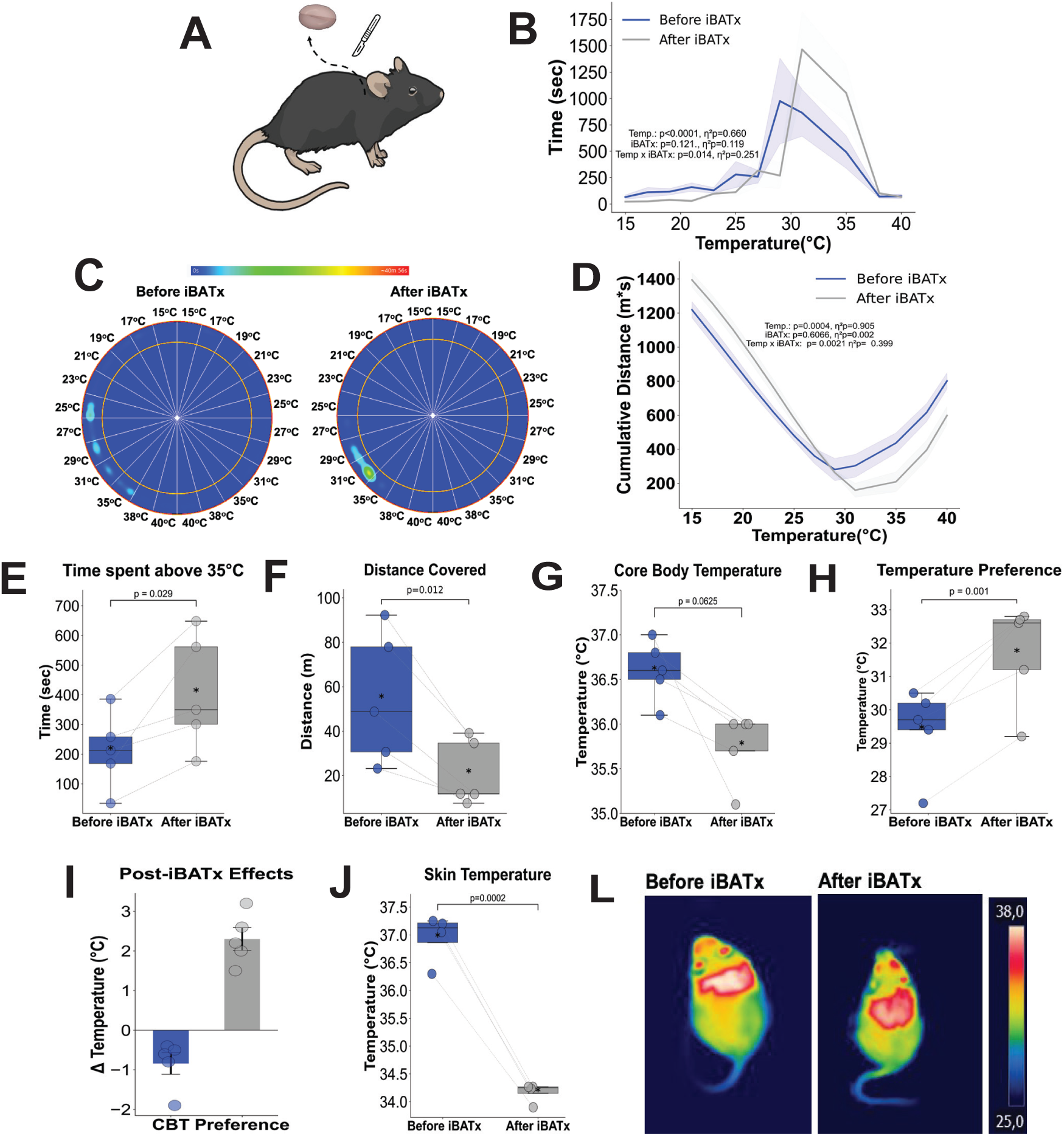
Interscapular brown adipose tissue lipectomy (iBATx) decreases core body temperature and shifts thermal preference toward warmer zones in mice. Eight-week-old male C57BL/6 mice (n = 5) were evaluated in a TGR system before and after 7 days of iBATx intervention. (A) Schematic of iBATx surgical protocol. (B) Time spent per temperature zone within the TGR system during a 60-minute observation period. (C) Representative thermal occupancy heatmap depicting murine locomotor patterns in the TGR. (D) Cumulative distance from the zones across experimental groups. (E) Time spent in temperature zones above 35°C within 60-minute interval. (F) Locomotor activity profiles during TGR assessment. (G) Rectal temperature measurements before and after intervention. (H) Mean of PT within the TGR system. (I) Comparative analysis of after iBATx Δ changes in CBT and PT. (J) Skin temperature proximal to BAT depots. (L) Representative infrared thermograms of BAT-associated regions. Data are presented as mean ± SEM; p < 0.05 (α = 5%). Statistical analysis: Two-way repeated-measures ANOVA (panels B-D); paired t-test (panels E-J).

### Genetic deficiency of UCP1 drives a warm-seeking behavior

Uncoupling protein 1(UCP-1) is one of the most studied and relevant mediators of thermogenesis in BAT (Aquila *et al*, 1985; Chouchani *et al*, 2019; Keipert *et al*, 2017; Fedorenko *et al*, 2012). Here, we investigated whether the lower thermogenic capacity caused by UCP-1 deletion is reflected by differences in temperature preference. Indeed, compared to WT littermate controls, UCP-1 deficient young male mice exhibited a shift to warmer zones in TGR analysis (<Figures 5B<). Representative heat map images of the mice running through the temperature zones and cumulative distance plot also corroborate the warm seeking phenotype of UCP1 deficient mice (<Figure 5C and D<). Additionally, UCP-1 KO mice exhibited higher warm tolerance as they stayed longer in temperatures above 35°C (<Figure 5E<). Interestingly, UCP-1 KO mice have lower mobility than WT mice, reflecting their need to warm up at high temperature zones (<Figure 5F<). We confirmed the reduction in rectal temperature in UCP-1 KO mice (<Figure 5G<), in contrast with increases in PT compared to WT mice (<Figure 5H<). Similar to the iBATx mice group, the PT variations observed in UCP-1 deficiency were greater than the rectal temperature (<Figure 5I<). Thermographic images did not show differences in superficial temperature between the WT and UCP-1 KO groups (<Figure 5J and L<). Altogether, these experiments indicate that either murine models of thermogenic gain or loss of function are adequately assessed by TGR method.

**Figure 5:**
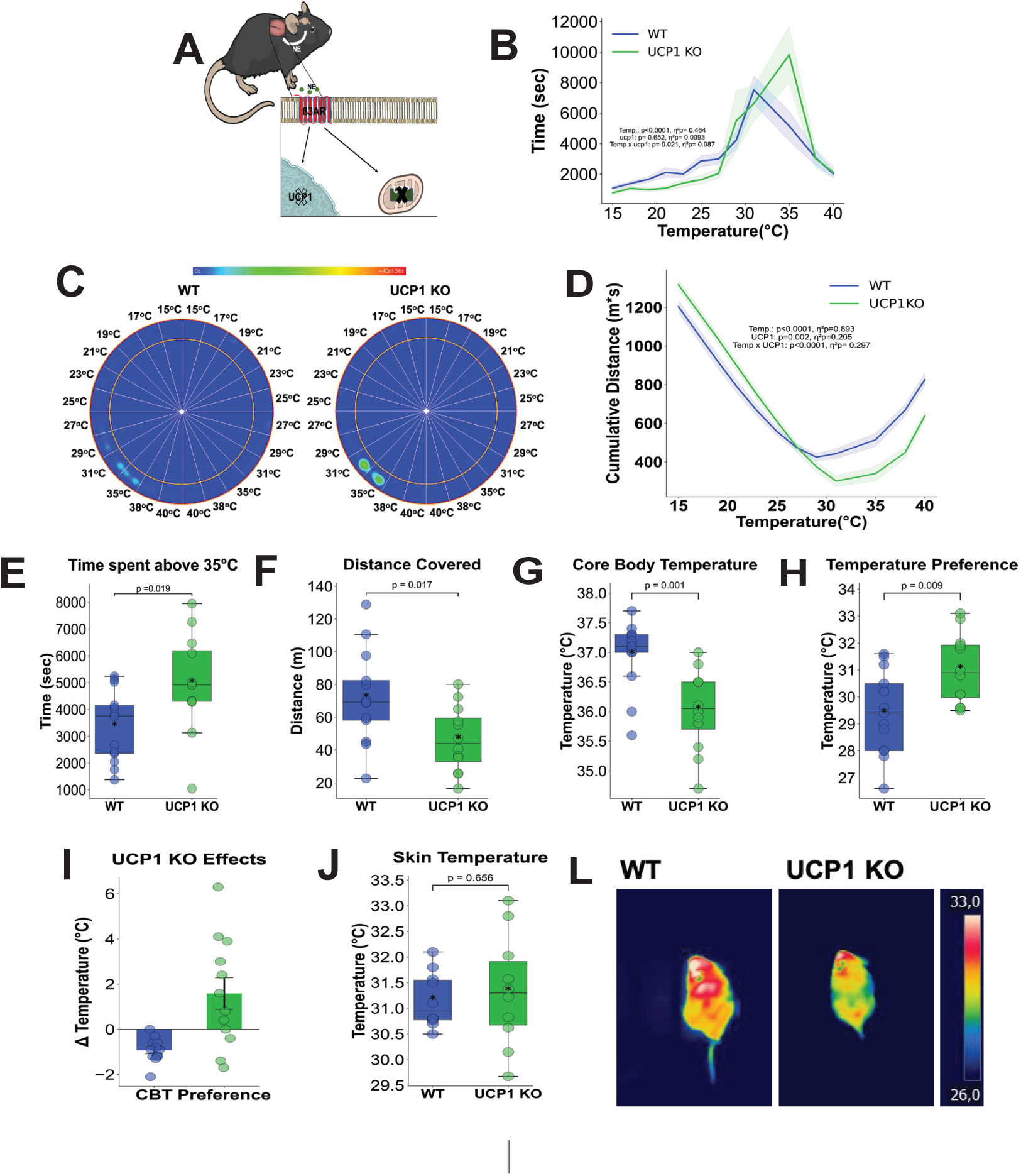
UCP1 genetic deficiency modifies thermal preference in mice. Eight-week-old male UCP1 knockout mice (n = 13) and WT littermate controls (n = 12) were sequentially evaluated in a TGR system. (A) Schematic of the experimental design comparing UCP1 KO and WT cohorts. (B) Time spent per temperature zone within the TGR system during a 60- minute observation period. (C) Representative thermal occupancy heatmap depicting murine locomotor patterns in the TGR. (D) Cumulative distance from the zones across genotypes. (E) Time spent in temperature zones above 35°C in a 60-minute interval. (F) Locomotor activity profiles during TGR assessment. (G) RT measurements. (H) Mean PT within the TGR system. (I) Comparative analysis of Δ differences in CBT and PT between WT and UCP1 KO mice. (J) Skin temperature proximal to brown adipose tissue (BAT) depots. (L) Representative infrared thermograms of BAT-associated regions. Data are presented as mean ± SEM; p < 0.05 (α = 5%). Statistical analysis: Two-way mixed-design ANOVA with between-subjects factor (panels B-D); unpaired t-test (panels E-J).

### Cold exposure renders a warm-seeking phenotype in TGR

Endothermy is a crucial feature of mammalian organisms that allows for the maintenance of the body’s physiological functions and biochemical reactions regardless of changes in environmental temperature. Endotherms regulate metabolic heat production through both shivering (ST) and non-shivering thermogenesis (NST) (Cannon & Nedergaard, 2011; Ivanov, 2006). During the initial phase of cold exposure, shivering predominates as a thermoregulatory mechanism until brown adipose tissue becomes fully activated. Once this occurs, NST takes over as the primary mechanism for maintaining thermal homeostasis (Cannon & Nedergaard, 2004; Cypess *et al*, 2009; Sponton *et al*, 2022). Although the NST mechanisms can keep CBT within a proper range for survival, this temperature usually stabilizes at a lower range in comparison to animals kept at room temperature (Cannon & Nedergaard, 2011; Keipert *et al*, 2017). We moved on to investigate whether cold exposed mice would display a thermogenic or a warm seeking phenotype at the TGR system (Figure 6A). Despite the elevated NST, cold-exposed mice displayed a preference for warmer zones, as shown by the absolute time spent in the zone, which is also illustrated by the heat map (Figure 6B and 6C). Cold-exposed mice exhibited an increase in heat tolerance, opting for temperatures and accumulating in zones above 35°C (Figure 6D and Figure 6E). Additionally, we found that cold-challenged mice exhibited a reduction in mobility within the TGR (Figure 6F), which presumably reflects the attempt to avoid heat loss. These TGR data indicated that despite the elevated thermogenic capacity, cold adapted mice prioritize restoring its CBT in a warmer surface. In line with these data, we observed that cold exposure reduced the average CBT by 1.5°C (Figure 6G and Supplementary Figure 1) and caused an elevation in PT of about 3°C (Figure 6H and Figure 6I). Interestingly, we also have employed a longer duration of cold exposure (21 days) and observed comparable warm-seeking behavior as demonstrated for the 4 days (Supplementary Figure 1). IR thermography analysis data confirmed the reduction in animal skin temperature, which is in agreement with the CBT and PT data (Figure 6J and Figure 6L). Together, these data indicate TGR is also a sensitive tool for evaluating fluctuations in endothermic behavior (warm seeking) associated with changes in environmental temperature.

**Figure 6:**
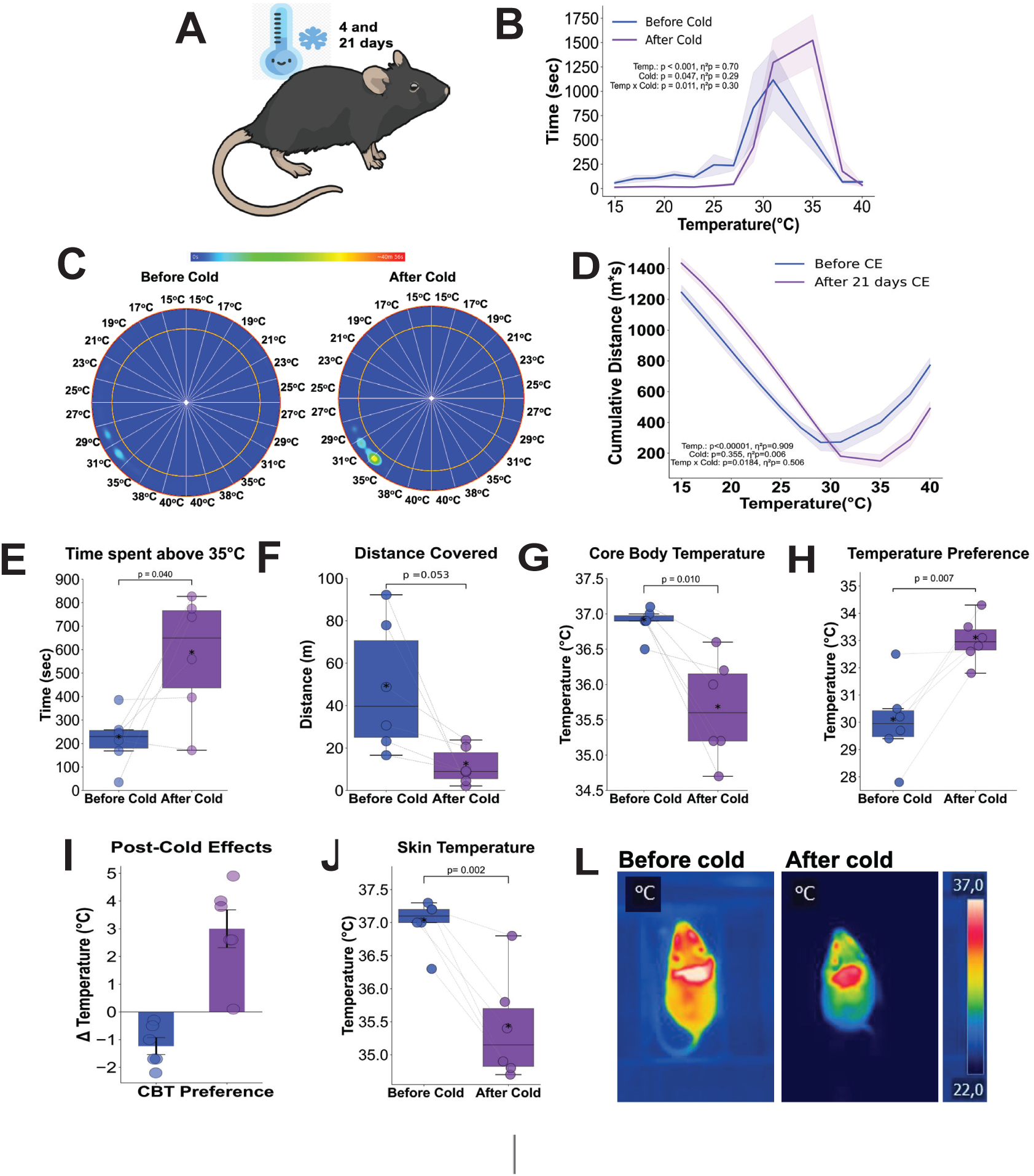
Cold acclimation elevates preferred temperature in mice. Eight-week-old male C57BL/6 mice (n = 6) were evaluated in a TGR system before and after 4 days cold acclimation (4°C). (A) Schematic of cold exposure experimental design. (B) Time spent per temperature zone within the TGR system during a 60-minute observation period. (C) Representative thermal occupancy heatmap depicting murine locomotor patterns in the TGR. (D) Cumulative distance from the coldest zone across experimental groups. (E) Time spent in temperature zones above 35°C within 60-minute interval. (F) Locomotor activity profiles during TGR assessment. (G) Rectal temperature measurements before and after acclimation. (H) Mean PT within the TGR system. (I) Comparative analysis of Δ changes in CBT and PT following cold acclimation. (J) Skin temperature proximal to BAT depots. (L) Representative infrared thermograms of BAT-associated regions. Data are presented as mean ± SEM; p < 0.05 (α = 5%). Statistical analysis: Two-way repeated-measures ANOVA (panels B, C, E); paired t-test (panels F–L).

### TGR analysis has proven to be a sensitive method for evaluating thermogenesis-associated behavior

Next, we examined the correlation between CBT measured via rectal temperature and PT assessed through TGR across all groups following gain or loss of function treatment. We observed a strong negative correlation between CB and PT values, indicating a highly significant association between both variables (Figure 7A). We also conducted a regression coefficient analysis on the changes observed in CBT and PT across all groups: CL injection, HFD, iBAT lipectomy, UCP1KO, and cold exposure mice. Interestingly, in all groups, PT values exhibited a greater degree of regression coefficient compared to CBT (Figure 7B), suggesting that PT are more sensitive to changes in thermogenesis than CBT. Indeed, PT depends on CBT fluctuations under metabolic interventions, as mice with BAT thermogenesis gain-of-function intervention seek cooler areas and exhibit lower PT, while thermogenesis loss-of-function mice seek warmer areas with an increase in PT (Figure 7C).

**Figure 7:**
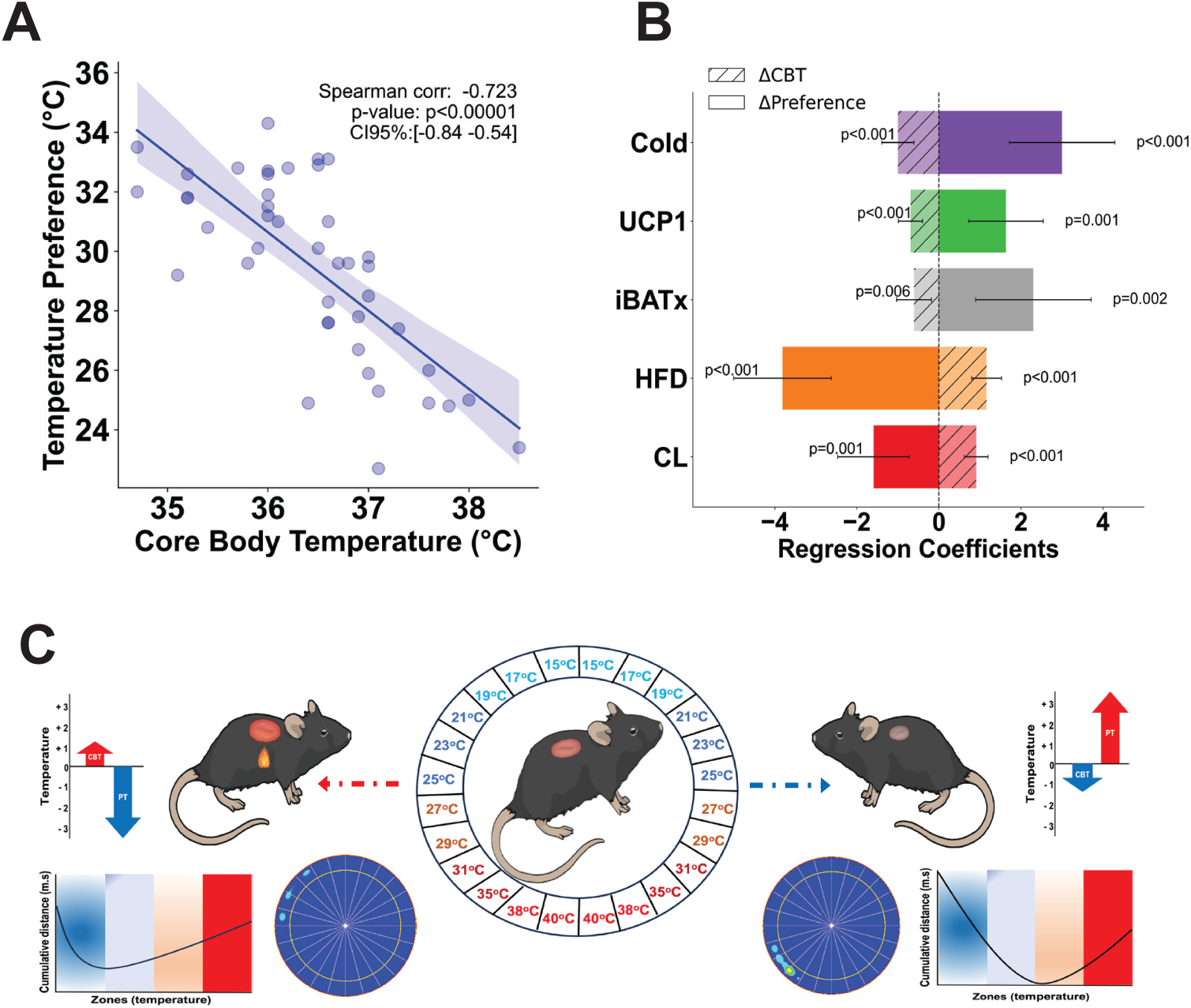
Temperature-associated behavioral analysis via TGR system demonstrates efficacy in evaluating thermogenesis for metabolic studies. Summary of experimental findings:(A) Spearman correlation analysis between core body temperature and preferred temperature across all treatment groups. (B) Multiple regression coefficients derived from treatment groups, comparing CBT (measured via rectal temperature) and PT (assessed via TGR). (C) Schematic representation of thermogenesis modulation outcomes: mice subjected to BAT thermogenesis gain-of-function interventions exhibit preferential occupancy of cooler zones with reduced PT, whereas BAT thermogenesis loss-of-function models display increased PT and prolonged occupancy in warmer regions.

## DISCUSSION

Developing new methods to assess body temperature in metabolic and thermogenesis research is essential for improving data reliability in the field. Metabolic interventions, such as alterations in food intake, energy expenditure, or thermal stress, are often reflected in temperature changes, highlighting the need for precise measurement techniques (Mei *et al*, 2018; Hymczak *et al*, 2021; Meyer *et al*, 2017). Since its development, the TGR system has been extensively used to study nociception, sensory nerve function, and thermal (Lei *et al*, 2023; Sasajima *et al*, 2022; Touska *et al*, 2016). While its primary applications have been in these fields, the potential for broader use in metabolic research and thermoregulation has remained an area for further exploration. Here, we demonstrate that the TGR method allows for the use of quantitative behavioral parameters that reflect the metabolic heat production capacity of the mice. We also show that such a system is adequate for the assessment of thermoregulatory state in either pro-thermogenic or thermo-incompetent mouse phenotypes.

As our data suggest, a key advantage of TGR is its high sensitivity to small fluctuations in heat flow, making it particularly useful for assessing thermogenesis in mice with a mild metabolic phenotype, whereas traditional methods may lack the resolution to detect minor thermogenic changes (Clement *et al*, 1989; Garami *et al*, 2011; Mei *et al*, 2018; Meyer *et al*, 2017). Another strength of TGR is its real-time, continuous monitoring capability, which allows us to have a thermogenic profile over time while rectal temperature and IR thermography only allow for intermittent (Mei *et al*, 2018; Touska *et al*, 2016). This feature is particularly valuable for studying dynamic metabolic adaptations, such as acute responses to environmental temperature changes or pharmacological interventions. Furthermore, TGR’s non-invasive nature represents a significant advantage over thermometry techniques that capture core temperature, which involve probe insertion in the rectum or through surgery. TGR system preserves the physiological integrity of the mice and minimizes potential artifacts caused by physical interference. Previous linear temperature gradient assays have utilized plexiglass tubes with copper coils or conductive floors to create gradients for measuring preferred temperatures in foot pads (Carlisle *et al*, 1999; Gordon *et al*, 2001, 2000; Dhaka et al., 2007). Those systems required observer intervention and were further limited by the tendency of mice to seek refuge in the corners of rectangular environments. In contrast, the automated TGR system improves upon these linear systems by eliminating these drawbacks, offering a more reliable and unbiased method for assessing thermotaxis.

Despite these advantages, TGR also has limitations. One notable constraint is the impossibility of integrating it with indirect calorimetry, a widely used method for assessing energy expenditure that often incorporates thermal probes (Hymczak *et al*, 2021; Ono-Moore *et al*, 2020). This makes it challenging to directly correlate heat production with metabolic rate measurements in a single experimental setup. Moreover, animals are analyzed individually, with each session lasting approximately one hour, whereas the inserted probe approach allows for the simultaneous recording of several animals. Rectal and skin temperature measurements are also quicker to take than TGR (Touska *et al*, 2016; Mei *et al*, 2018; Meyer *et al*, 2017; Zethof *et al*, 1994). Interpreting TGR data in cold-exposed mice requires careful consideration. When challenged by cold, mice exhibit increased heat production through thermogenesis; however, their overall body temperature remains low due to external cooling (Chau *et al*, 2024; Sato *et al*, 2015). Our findings suggest that under such conditions, mice prioritize warmth-seeking behavior to restore CBT rather than sustaining high thermogenic activity. This behavioral adaptation must be accounted for TGR analyses, as it represents an energy-efficient thermoregulatory strategy rather than a direct measure of thermogenic output. This aligns with the hierarchical organization of thermoregulation in mammals, where strategies are prioritized based on energy cost (Glass *et al*, 2021; Ivanov, 2006). Non-shivering thermogenesis is the most energy-demanding adaptive response, whereas seeking a warmer environment is a low-cost behavioral adjustment. Since cold-exposed mice experience lower core temperatures, they instinctively move toward warmer zones in the TGR system to quickly normalize their body temperature at a low energy cost, thereby reducing the need for prolonged thermogenic activation (Cannon & Nedergaard, 2011; Glass *et al*, 2021).

It is important to highlight that the TGR system provides an indirect, behavior-based assessment of thermogenesis rather than a direct measurement of body temperature, as other methods do. However, this behavioral approach offers valuable insights into thermoregulatory strategies, capturing how animals actively regulate their thermal environment. Our observations revealed that mice with both high and low thermogenic capacities tend to reduce movement, preferring to remain in a single zone, either to conserve heat in warmer zones or to minimize heat generation through activity. An exception to this pattern was observed in high-fat diet mice, which exhibited normal movement patterns. This behavior is likely a consequence of higher energy intake from a hypercaloric diet, necessitating greater energy expenditure to maintain metabolic balance (Ono-Moore *et al*, 2020). Notably, high-fat-fed mice exhibited a preference for moving through cooler zones, likely increasing their energy expenditure and stimulating heat production as a compensatory strategy to prevent excessive energy storage and mitigate the metabolic burden associated with a surplus caloric intake (Feldmann *et al*, 2009; Nedergaard & Cannon, 2022). These findings underscore the interaction between thermoregulatory behavior and physiological thermogenesis, demonstrating that behavioral adaptations play a crucial role in metabolic regulation. By integrating both aspects, the TGR system provides a more comprehensive perspective on energy balance and thermoregulation in different metabolic states.

In conclusion, the TGR system represents a powerful, sensitive, unbiased, and non-invasive tool for studying thermogenesis and metabolic regulation. While it provides superior sensitivity and real-time monitoring compared to traditional thermometry, its integration with other metabolic assessment techniques remains an area for future development. Moreover, interpreting data in variable environmental conditions requires careful analysis of both thermogenic responses and behavioral adaptations. Nonetheless, TGR has strong potential to enhance our understanding of metabolic thermoregulation-associated behaviors across a number of research applications.

