## Supplementary Figure 1 for "Non-invasive and unbiased assessment of thermogenesis in mice through thermal gradient ring"

### De Melo et al., 2025. Supplementary Figure:

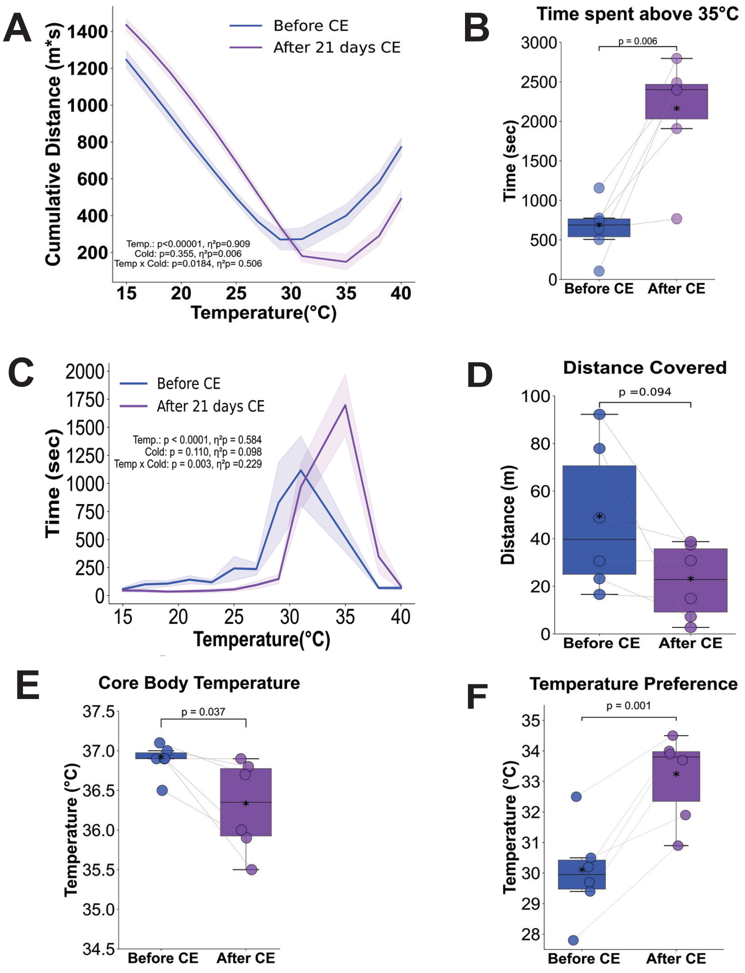


**Supplementary Figure 1: Long-term cold exposure elevates preferred temperature in mice.** Eight-week-old male C57BL/6 mice ($n = 6$) were evaluated in a TGR system pre- and post-21-day cold acclimation (4°C). (A) Cumulative distance from the coldest zone across experimental groups. (B) Time spent in temperature zones >35°C within a 60-minute period. (C) Total time spent per temperature zone within the TGR apparatus. (D) Locomotor activity profiles during TGR assessment. (E) Rectal temperature measurements before and after acclimation. (F) Mean PT within the TGR system. Two factors ANOVA for repeated measurements (A and C), Paired T-test (B-F), $\alpha = 5\%$.
